# Field-based body temperatures reveal behavioral thermoregulation strategies of the Atlantic marsh fiddler crab *Minuca pugnax*

**DOI:** 10.1101/2020.07.03.187229

**Authors:** Sarah Hews, Zahkeyah Allen, Adrienne Baxter, Jacquline Rich, Zahida Sheik, Kayla Taylor, Jenny Wu, Heidi Zakoul, Renae Brodie

**Author notes:** These authors contributed equally to this work (Sarah Hews, Renae Brodie).

## Abstract

Behavioral thermoregulation is an important defense against the negative impacts of climate change for ectotherms. In this study we examined the use of burrows by a common intertidal crab, *Minuca pugnax*, to control body temperature. To understand how body temperatures respond to changes in the surface temperature and explore how efficiently crabs exploit the cooling potential of burrows to thermoregulate, we measured body, surface, and burrow temperature data during low tide on Sapelo Island, GA in March, May, August, and September of 2019. We found that an increase in 1°C in the surface temperature led to a 0.70-0.71°C increase in body temperature for females and an increase in 0.75-0.77°C in body temperature for males. Body temperatures of small females were 0.3°C warmer than large females for the same surface temperature. Female crabs used burrows more efficiently for thermoregulation compared to the males. Specifically, an increase of 1°C in the cooling capacity (the difference between the burrow temperature and the surface temperature) led to an increase of 0.42-0.50°C for females and 0.34-0.35°C for males in the thermoregulation capacity (the difference between body temperature and surface temperature). The body temperature that crabs began to use burrows to thermoregulate was estimated to be around 24°C, which is far below the critical body temperatures that could lead to death. Many crabs experience body temperatures of 24°C early in the reproductive season, several months before the hottest days of the year. Because the use of burrows involves fitness trade-offs, these results suggest that warming temperatures could begin to impact crabs far earlier in the year than expected.

## Introduction

Warmer than average days and heat waves are becoming more frequent with climate change ([1]). For ectothermic organisms that thermoregulate to keep their body temperatures close to a thermal optimum (*T*_*o*_), the physiological responses to elevated environmental temperatures are understood. Excessively warm days exert deleterious impacts on energy and water budgets ([2]; [3]; [4]), and more extreme environmental temperatures lead to anaerobic respiration, compromising tissue function ([5]; [6]; [7]). Deviations from *T*_*o*_, including those that surpass critical thermal limits for short periods, can be survivable because organisms have compensatory mechanisms. For example, at a cellular level organisms can up-regulate genes associated with repair processes ([8]; [9]; [10]), and they can engage in behaviors that increase tissue oxygenation once a crisis has passed ([11]). However, compared to endotherms, ectotherms have less control over internal processes that can generate or redistribute heat within their bodies and rely heavily on behavioral mechanisms to regulate body temperatures ([12]; [13]). Under the likely scenario of continued warming, behavioral thermoregulation will be a critical means for ectotherms to avoid extreme temperatures. For this reason, the efficiency with which they can use the surrounding environment to adjust their body temperatures and the costs associated with these behaviors are important determinants of vulnerability.

As a highly active, intertidal forager, the fiddler crab is an excellent model system in which to investigate the impacts of environmental temperatures on thermoregulatory behaviors. During daytime low tides, droves of foraging individuals occupy exposed mud and sand flats where they feed on microflora and fauna ([13]; [14]; [15]). Fiddler crabs, like other ectotherms, employ a variety of physiological, morphological, and behavioral strategies to thermoregulate during these periods of exposure. Evaporative cooling from a wetted body is possible on windy and less humid days ([16]; [17]), and some species can readily change the distribution of chromatophores on their cuticles to increase reflectance ([18]; [19]). On hot days, fiddler crabs orient their bodies to minimize the surface area experiencing direct exposure from the sun ([17]) and males radiate heat from their enlarged claw ([20]) that is also an ornament and weapon used in courtship contests ([21]; [22]). As mobile ectotherms, they have the option to retreat to cooler microhabitats, including shade ([23]; [17]), and especially the burrow ([17]; [16]; [23]).

For fiddler crabs, offloading heat in a burrow that is cooler than the surface is a highly effective thermoregulatory strategy ([17]; [24]). Burrow temperatures decline exponentially with depth and burrows maintain a more stable temperature profile compared to the surface ([16]). Semi-permanent burrows extending 10-60 cm ([25]; [26]; [27]) into the substratum can exceed densities of 100 burrows/m2 in areas occupied by fiddler crab colonies (e.g., [28]; [29]). Individual crabs can claim a burrow, modify it, and use it as a place to hide from predators ([30]; [31]), to court, to mate, to incubate embryos ([26]), and use it for thermoregulation. Individuals that are foraging rather than defending a burrow are rarely more than a few body lengths from a burrow and can access unoccupied or poorly defended burrows as they travel across the substratum (pers obs). The cooling capacity and abundance of burrows may explain why researchers report finding fiddler crabs active when surface temperatures exceed their critical thermal limits ([32]; [23], pers. obs.).

It is now recognized that ectotherm survival in warming habitats may depend on their ability to exploit microclimates to manage body temperatures ([33]; [34]). Here, we used the Atlantic marsh fiddler crab *Minuca pugnax*, a temperate species with a range from New Hampshire to northern Florida USA ([35]), as a model system to investigate the relationship between *T*_*b*_ and the temperature of surface and burrow sediments (hereafter referred to as surface and burrow temperatures). *T*_*b*_ is influenced by heat exchange between the organism and the environment through radiation from the sun and other objects, as well as convection and conduction ([36]). We measured sediment temperatures because fiddler crabs, like many other errant marine intertidal invertebrates, have wet bodies that are in nearly constant contact with wet sediment, making thermal flux between the surface, burrow and body an important determinant of *T*_*b*_. Finally, we explored the usefulness and limits of burrows for thermoregulation. In this study, we employed several parameters that are commonly used in thermal ecology research, and introduced two new parameters, *T*_*reg*_ and *E*_*B*_ (see Table 1 for definitions of parameters measured or mentioned in this study). Specifically, we asked: (1) How is *T*_*b*_ impacted by surface temperatures and what is the range of *T*_*b*_ experienced during the active season (spring, summer and fall) near the southern end of the species range; (2) How efficiently can crabs exploit the cooling potential of burrows to thermoregulate, and how do changes in the cooling potential of the burrows impact thermoregulation (*E*_*B*_); and (3) At what *T*_*b*_ does a crab begin to use burrows to resist heat transfer from the environment (when does *T*_*b*_ = *T*_*reg*_)?

**Table 1.** Definitions of parameters measured or mentioned in this study. All parameters are body temperatures.

| Parameter | Description |
| --- | --- |
| $T_b$ | Body temperature. For this study, all $T_b$ measurements were from field-based individuals |
| $T_o$ | The body temperature at which performance is optimized (Huey and Kingsolver 1989; Gilchrist 1995; Angilletta et al. 2002). |
| $T_{set}$ | Set point or target $T_b$. The preferred body temperature chosen by an ectotherm on a thermal gradient (Huey et al. 1989; Huey 1991; Hertz et al. 1993). |
| $T_{reg}$ | The lowest body temperature at which an ectotherm starts to thermoregulate using the burrows to avoid overheating. |
| $T_e$ | Equilibrium or operative body temperature. Body temperature of a non-thermoregulating control organism (the null model), measured directly from physical models (biomimics) or estimated mathematically (Heath 1964; Huey 1974; Roughgarden et al. 1981). In this study, a modified $T_e(B)$ was calculated for free ranging crabs that could not use burrows for thermoregulation, but potentially could have used other means. |
| $E_B$ | Burrow use efficiency. Change in the thermoregulation efficiency (body temperature minus surface temperature) due to a 1°C change in the cooling capacity (burrow temperature minus surface temperature) of the burrow. In this study we measured the burrow use efficiency of *M. pugnax* exploiting burrows. |
| $CT_{max}$ | The maximum $T_b$ at which performance is possible. For fiddler crabs this has been measured as the $T_b$ at which they lose the righting response (e.g. [23]; [37]) |

## Materials and methods

### Field based body temperatures (*T*_*b*_)

The body temperatures (*T*_*b*_) of surface active male and female *Minuca pugnax* were measured on ten different days between May and October 2019 on Sapelo Island, Georgia, USA at Shell Hammock (31.399832,-81.287417) and lighthouse road (31.390724, -81.285953), using a Physitemp Instruments portable temperature monitor (PTM1) with a Type T needle microprobe (MT-29). Individual crabs of around 1 cm carapace width or larger were picked up from the surface opportunistically and the needle probe was inserted between the 2nd and 3rd walking leg into the gill chamber for a temperature reading (0.1°). Following the *T*_*b*_ measurement, the carapace width (cw) was measured with a digital caliper (0.1 mm) and the crab was released. The same microprobe was used to measure the surface temperature at the location where each individual crab was caught, and every 10-30 minutes during the collection period a temperature measurement was taken from the bottom of a nearby artificial burrow made from a 2.5cm wide pvc pipe, extending 30 cm into the substratum. This reference depth was within the range that crabs can access and ensured that the measurement captured the coolest burrow microclimate available at that time and place, as burrow temperatures decline exponentially with depth, with most of that change occurring between the surface and a depth of 15 cm ([38]). The artificial burrow was moved frequently to keep it close to areas where crabs were being measured.

Males and females of the same carapace width are not comparable because of the male’s enlarged sexually selected claw. For this reason, in most of the statistical analyses, size was treated as a dichotomous categorical variable divided at the midpoint in the range of sizes collected for each sex. Four classes of crabs were created: small females (9-12.9 mm cw), large females (13-17mm cw), small males (10-14.9 mm cw), and large males (15-20 mm cw).

To determine if there were differences in *T*_*b*_ between the four categories of crab, we used a Kruskal-Wallis H test followed by pairwise comparisons with a Dunn’s (1964) procedure and a Bonferonni correction for multiple comparisons. An ANOVA was not used because the assumption of normality was not met after multiple transformations were tried. The four sampling months, March, May, August and October 2019 were analyzed separately.

Using the field based *T*_*b*_ and surface temperature (*S*) measurements described above, we calculated the difference between the body and surface temperature, *T*_*b*_ − *S*, for each crab, where a positive value indicated that the crab was warmer than the surface and a negative value indicated that it was cooler. Separate regressions for the four categories of crab, with surface temperature as the independent variable and *T*_*b*_ − *S* as the dependent variable, were plotted and used to investigate crab abilities to maintain body temperatures different from the surface, including the specific *T*_*b*_ at which crabs began to use burrows to cool themselves (the *T*_*reg*_ estimation described below). For each sex, we investigated the impact of size with an ANCOVA, where *T*_*b*_ − *S* was the continuous dependent variable, crab size was the categorical independent variable, and surface temperature was the continuous covariate. We tested for the homogeneity of regression slopes by investigating the interaction between surface temperature and crab type in the GLM. For females, untransformed values of the dependent variable were used and the model met assumptions of linearity, homogeneity of regression slopes, normality, homoscedasticity, homogeneity of variances, and there were no outliers. While most of the model assumptions were met for males, the residuals for large males were not normally distributed, the homogeneity of variances was violated, and there were four outliers. While a log10 transformation addressed the problem with homogeneity of variances and adjusted three of the four outliers, no transformation was found that could normalize the residuals. For this reason, large and small males were not compared to each other in a statistical analysis.

### Estimating the burrow use efficiency (*E*_*B*_) and an estimate for the difference between the operative body temperature in the absence of burrows (*T*_*e*_(*B*)) and the surface (*S*)

The burrow use efficiency, *E*_*B*_, was determined by finding the rate of change of the relationship between two newly defined terms: (1) the cooling capacity of the burrow as the potential of the burrow to cool the crab, and (2) the thermoregulation capacity as the ability of the crab to use the burrow and other means to regulate body temperature. The larger the cooling capacity, the greater the potential for the burrow to keep the crab cool when the surface temperature is too warm. (Theoretically, the burrow could also be used to heat the crab when the burrow is warmer than the surface.) The greater the difference between crab and surface temperature, the larger the crab’s thermoregulation capacity or the degree to which it was managing its body temperature through behavioral, physiological, and morphological mechanisms.

The cooling capacity of the burrow, represented by the *x* variable, is the difference between the burrow temperature and the surface temperature, *x* = *B* − *S*. The thermoregulation capacity of the crab, represented by the *y* variable, is the difference between the crab body temperature and the surface temperature, *y* = *T*_*b*_ − *S*.

We represent the relationship between the cooling/heating capacity of the burrow and the thermoregulation capacity of the crab in a coordinate plane (Fig 1A). We call this representation of the relationship the thermoregulation axes. The blue thermoregulatory zone corresponds to *x, y* < 0, where both the burrow and the crab are cooler than the surface. Theoretically, this zone represents the crab using the burrow to thermoregulate when the surface is too warm. The red thermoregulatory zone corresponds to *x, y >* 0, where the burrow and the crab are warmer than the surface, and represents the crab using the burrow to warm itself through thermoregulation when the surface is too cold.

**Fig 1.**
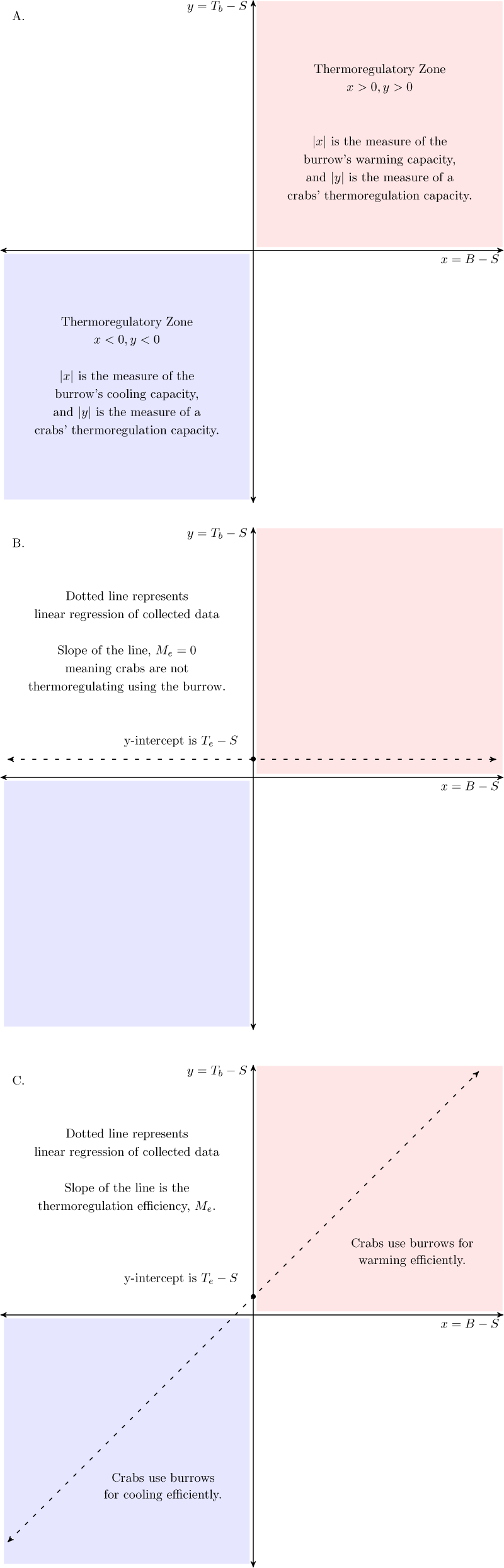
The thermoregulation axis. The graph of the thermoregulation capacity (*T*_*b*_ − *S*) against the cooling capacity (*B* − *S*) of the burrow. A: Describes the thermoregulatory zones. B: The crabs are not thermoregulating using the burrow. C: The crabs are using the burrows extremely efficiently for cooling.

In the scenario where the crabs do not use the burrow to thermoregulate, then data collected of crab, burrow, and surface temperature would fall along a horizontal line (Fig 1B). If the crabs use the burrow to completely regulate their temperature, then data collected of crab, burrow, and surface temperature would fall along a line with slope 1 (Fig 1C). Both scenarios would have data with a positive y-intercept that corresponds to the difference between the surface temperature and the crab body temperature. When burrows are used to thermoregulate, the y-intercept marks the point where no thermoregulation with the burrow is occurring because burrow and surface temperatures are the same. We expect the y-intercept to be greater than zero because the thermal properties of their bodies cause crabs to be warmer than the surface when they are not thermoregulating. On a sunny day in August, 2019, we found that dead crabs and crabs filled with silicone (also dead) on an unshaded area of the marsh were 0.5-2.0°C warmer than the surface after 30 min (unpublished data for males).

The crab, burrow, and surface temperature data therefore falls on a line *y* = *E*_*B*_*x* + *b* on the thermoregulation axes where *E*_*B*_ is defined as the burrow use efficiency and *b* is how much warmer crabs are than the surface and set by material properties of the organism. *b* can also provide another estimate for the difference between the operative body temperature in the absence of burrows and the surface temperature, *b* = *T*_*e*_(*B*) − *S*.

The relationship between the cooling/heating capacity of the burrow and the thermoregulation capacity (the thermoregulation axes) were investigated for females using an ANCOVA analysis, where the thermoregulation capacity of the crab (*T*_*b*_ − *S*) was the continuous dependent variable, crab size (large and small) was the categorical independent variable, and the cooling capacity of the burrow (*B* − *S*) was the continuous covariate. A square root transformation of the dependent variable was used for females to meet the assumption of normality. The residuals for large males were not normally distributed and this could not be fixed with transformations. For this reason, large and small males were not compared in an ANCOVA analysis.

### Estimating the temperature that the crab begins to thermoregulate (*T*_*reg*_) due to the burrow

An estimate for *T*_*reg*_, the temperature that the crab begins to thermoregulate, is calculated by using the linear regression of *T*_*b*_ − *S* vs. *S* to find the surface temperature when *T*_*b*_ − *S* = *T*_*e*_(*B*) − *S* (this is the surface temperature when the crab begins to thermoregulate, 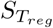), and then using the estimate of *T*_*e*_(*B*) − *S* to calculate the body temperature (*T*_*reg*_ = *ST*_*reg*_ + (*T*_*e*_(*B*) − *S*)) (Fig 2). As an estimate for *T*_*e*_(*B*) − *S*, we used the y-intercept from the thermoregulation axes for each crab category, which identifies how much warmer *T*_*b*_ is than the surface when burrows cannot be used for thermoregulation.

**Fig 2.**
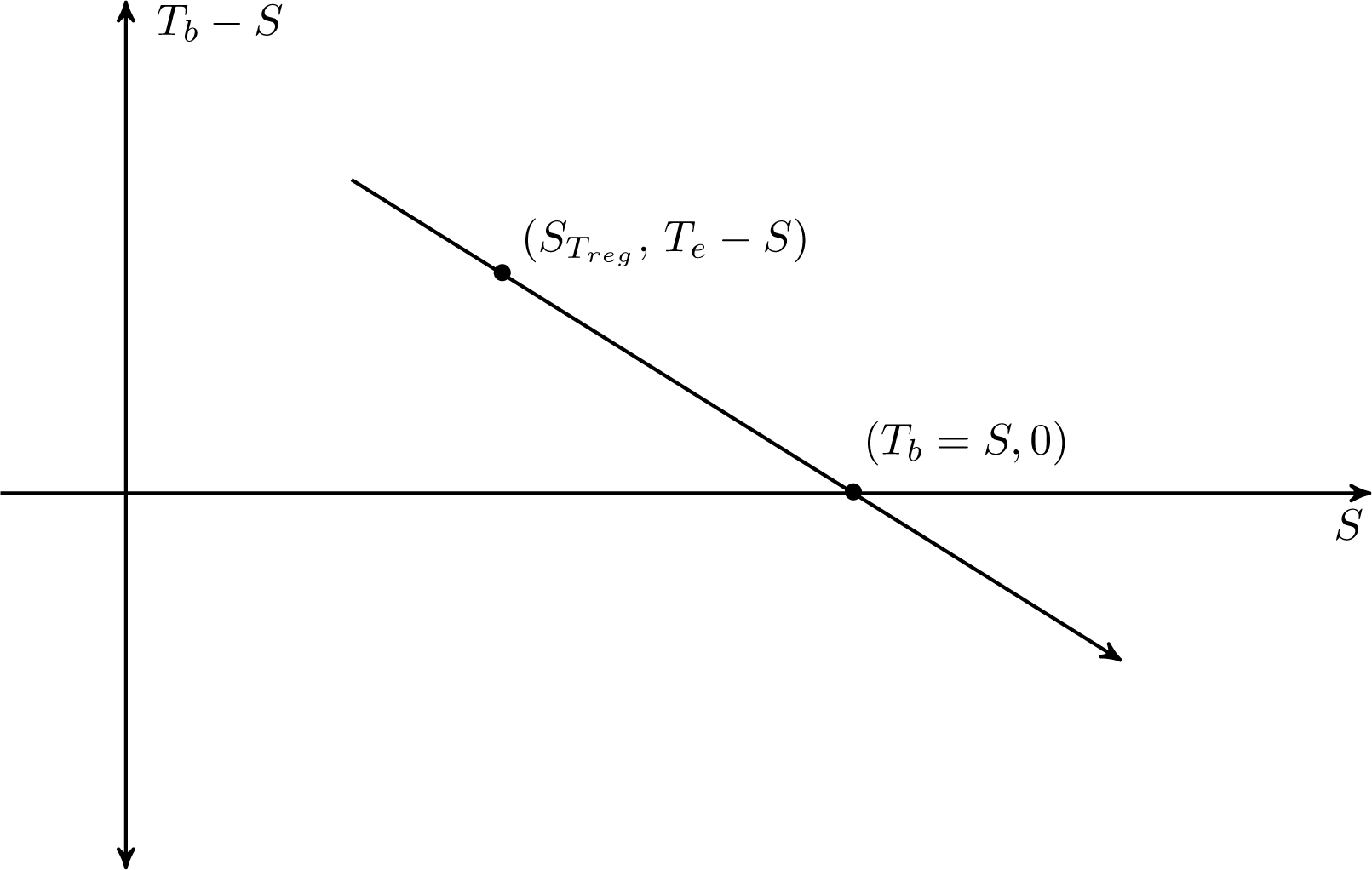
Method to estimate *T*_*reg*_, which is the lowest body temperature the crab begins to thermoregulate using the burrows. *T*_*b*_ is the crab body temperature, *S* is the surface temperature, *T*_*e*_(*B*) is the operative body temperature when crabs can’t use burrows, and 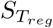 is the lowest surface temperature when the crab begins to thermoregulate. 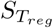 is found by establishing the linear relationship between *T*_*b*_ − *S* and *S* and determining the surface temperature where *T*_*b*_ − *S* = *T*_*e*_(*B*) − *S* (which is approximately 1.4°), or *T*_*b*_ = *T*_*e*_(*B*). Then, 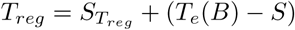. When *T*_*b*_ < *T*_*e*_, then thermoregulation must be occurring.

## Results

### Field based body temperatures (*T*_*b*_) during the active season, and the influence of size and surface temperature on thermoregulation capacity (*T*_*b*_ − *S*)

*T*_*b*_ measurements of 441 male (*n* = 252) and female (*n* = 189) *M. pugnax* were collected during low tide in 2019 on 29 and 30 March; 10, 11 and 13 May; 24, 26 and 28 August, and 12 and 13 October. Median *T*_*b*_s in March were 23.5°C (*n* = 22; small females), 23.4°C (*n* = 19; large females), 22.7°C (*n* = 38; small males), and 25.1°C (*n* = 38; large males) (Fig 3). There was an overall significant difference among *T*_*b*_ distributions in March (*H*(3) = 7.89, *p* = 0.048), where small and large males showed significantly different distributions from each other (Fig 3). In May, median *T*_*b*_s were 27.2°C (*n* = 19; small females), 27.3°C (*n* = 51; large females), 26.6°C (*n* = 48; small males) and 26.2°C (*n* = 29; large males) and there were no significant differences in *T*_*b*_ across groups (*H*(3) = 4.08, *p* = 0.253; Fig 3). Median *T*_*b*_ was highest in August at 34.5°C (*n* = 24; small females), 36.1°C (*n* = 22; large females), 33.8°C (*n* = 32; small males), and 32.8°C (*n* = 35; large males). There was an overall significant difference in *T*_*b*_ distributions in August (*H*(3) = 11.43, *p* = 0.01), with a significant difference between large males and large females (*p* = 0.006; Fig 3). In October, median *T*_*b*_s were 31.3°C (*n* = 5; small females), 30.5°C (*n* = 27; large females), 30.6°C (*n* = 9; small males), 31.1°C (*n* = 23; large males). No significant differences in *T*_*b*_ distributions were found for October (*H*(3) = 2.97, *p* = 0.413; Fig 3); however, the low sample sizes in two of the groups likely resulted in a low statistical power for this analysis.

**Fig 3.**
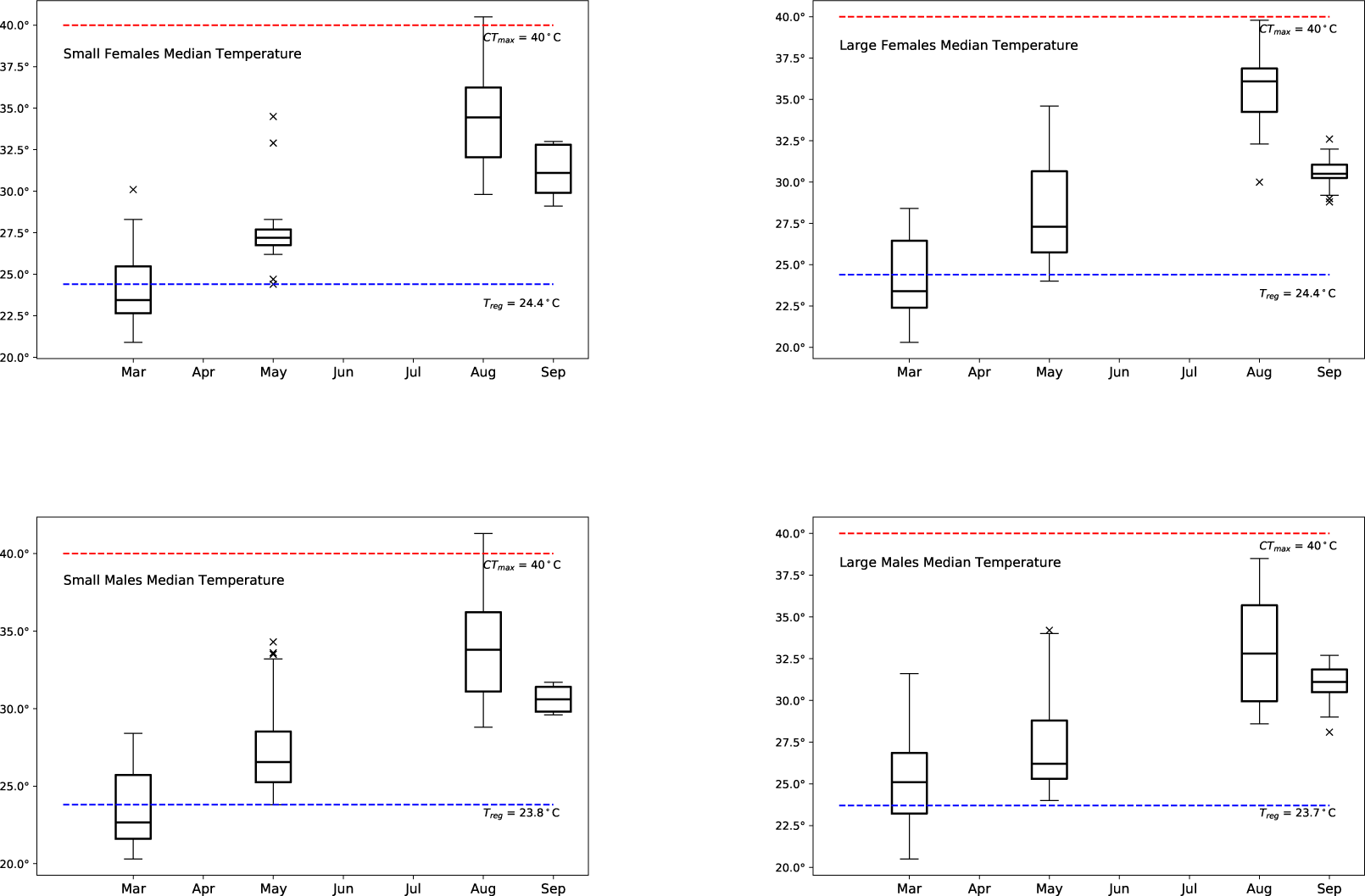
Box plots showing *T*_*b*_ for small females, large females, small males, and large males for March, May, August, and September. Red line represents *CT*_*max*_ and blue line represents estimates of *T*_*reg*_.

The relationship between the thermoregulation capacity (*T*_*b*_ − *S*) and the surface temperature (S) was determined to be linear for females and small males, with 79% (small females), 70% (large females), 61% (small males), and 42% (large males) of the thermoregulation capacity explained by the surface temperature (Fig 4).

**Fig 4.**
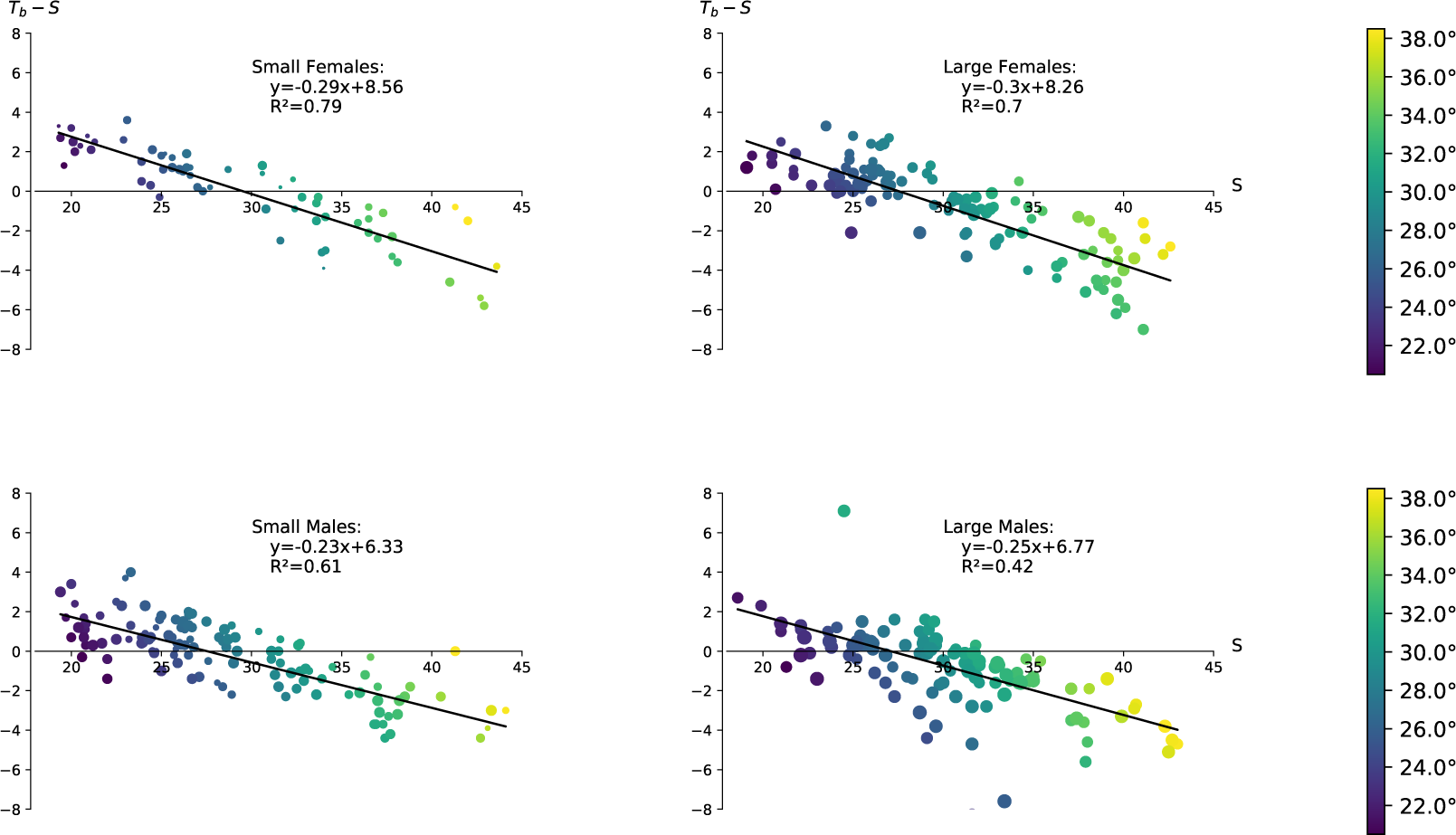
*T*_*b*_ − *S* against *S* and linear regressions for small females, large females, small males, and large males. Data was collected in March, May, August, and October, 2019.

For females, both the surface temperature (*F*_1,154_ = 454.2, *p* < 0.0005) and crab carapace width (*F*_1,154_ = 7.8, *p* = 0.006) were significant predictors of *T*_*b*_ − *S* and there were no interaction effects (*F*_1,154_ = .146, *p* = .703). The interpretation of these results is that for every 1°C increase in surface temperature, the thermoregulation capacity of females decreased by approximately 0.29°C - 0.30°C. An alternative interpretation is that for every increase in surface temperature of 1°C, *T*_*b*_ in females increased by 0.70°C-0.71°C. Carapace width being significant implies that the *T*_*b*_ of small females was 0.3°C warmer than large females for the same surface temperature (Fig 4). These results are congruent when carapace width is treated as a continuous variable.

Since the large male data were not normally distributed and could not be transformed to be so, we could not determine whether the difference between small males and large males was significant. The interpretation of the individual regressions for males is that for every 1°C increase in surface temperature, the thermoregulation capacity of males decreased by approximately 0.23°C -0.25°C. The alternative interpretation is that for every increase in surface temperature of 1°C, the *T*_*b*_ for males increased by 0.75°C-0.77°C. (Fig 4).

### Estimating the burrow use efficiency (*E*_*B*_) and an estimate for the difference between the operative body temperature in the absence of burrows (*T*_*e*_(*B*)) and the surface (*S*)

The relationship between the thermoregulation capacity (*T*_*b*_ − *S*), and the cooling capacity of the burrow (*B* − *S*) was determined to be linear for females and small males, with 68% (small females), 73% (large females), 61% (small males), and 50% (large males) of the thermoregulation capacity explained by the cooling capacity of the burrow (Fig 5).

**Fig 5.**
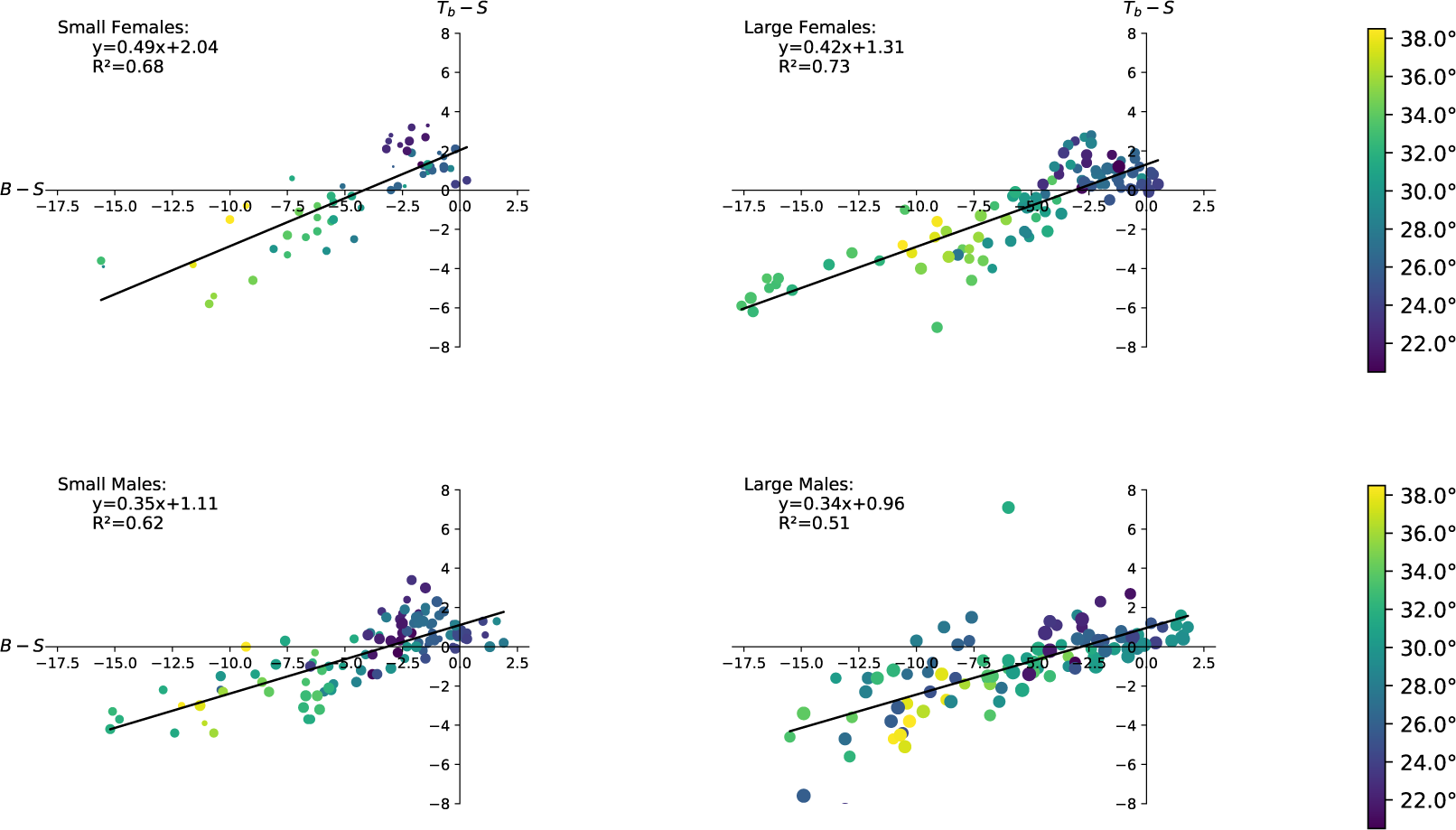
*T*_*b*_ − *S* against *B* − *S* and linear regressions for small females, large females, small males, and large males. Data was collected in March, May, August, and October, 2019.

For females, the cooling capacity (*F*_1,154_ = 330.6, *p* < 0.0005) was a significant predictor of the thermoregulation capacity and the crab carapace width (*F*_1,154_ = 3.8, *p* = 0.054) was a marginally significant predictor of the thermoregulation capacity. There were no interaction effects (*F*_1,154_ = 3.4, *p* = .069). This implies that the burrow use efficiency (*E*_*B*_) was not significantly different between small females (*E*_*B*_ = 0.5) and large females (*E*_*B*_ = 0.42), meaning that there was no significant difference in how thermoregulation capacity changes in response to changes in the cooling capacity of the burrow. For each 1° increase in the cooling capacity (meaning *B* − *S* decreases by 1°), females could increase their thermoregulation capacity by 0.42°C-0.50°C (meaning *T*_*b*_ − *S* decreases by 0.42°C-0.50°C). For a set cooling capacity, the thermoregulation capacity was 0.7°C greater for small females than large females. Alternatively, large females reduced their temperature by 0.7°C more than small females for a given cooling capacity and the same environmental conditions (same *B* and *S*). The y-intercept was 2.04°C for small females and 1.31°C for large females. This means that when the burrow temperature was equal to the surface temperature, *T*_*b*_ −*S* = *T*_*e*_(*B*) − *S*, the difference between the operative body temperature in the absence of burrows and the surface temperature was 2.04°C for small females and 1.31°C for large females (Fig 5). These results are congruent when carapace width is treated as a continuous variable.

Since the large male data were not normally distributed and could not be transformed to be so, we could not determine whether the difference between small males and large males was significant. For each 1°C increase in the cooling capacity (meaning *B* − *S* decreases by 1°C), males can increase their thermoregulation capacity by 0.34°C-0.35°C (meaning *T*_*b*_ − *S* decreases by 0.34°C-0.35°C). When the burrow temperature is equal to the surface temperature, *T*_*b*_ − *S* = *T*_*e*_(*B*) − *S*, the difference between the operative body temperature and the surface temperature for males was between 0.96°C-1.11°C (Fig 5).

### Estimating the temperature that the crab begins to thermoregulate (*T*_*reg*_) due to the burrow

Using the estimates for *T*_*e*_(*B*) − *S* (2.04°, small females; 1.31°, large females); 1.11°, small males; 0.96°, large males) (Fig 5), and the regression lines establishing the linear relationship between *T*_*b*_ − *S* and *S* (Fig 4), *T*_*reg*_ was calculated as 24.52° (small females), 24.48° (large females), 23.81° (small males), and 24.20° (large males).

## Discussion

Ectotherms like *Minuca pugnax*, will experience increased maintenance costs as air, water, and surface temperatures warm with climate change, necessitating compensatory responses that include behavioral thermoregulation. The critical thermal body temperatures (*CT*_*max*_) of ten different fiddler crab species lie between 40-43°C, with *M. pugnax* showing a *CT*_*max*_ of 40°C ([39]; [40]; [37]; [32];). This falls within the typical range for ectotherms, which across taxa, habitats, and latitudes, experience severe heat stress at *T*_*b*_s above 40°C with some exceptions ([41]; [42]; [43]; [33]). The median daytime *T*_*b*_s for *M. pugnax* was well below this during our August data collection period at 33°C and 36°C for large males and large females, respectively. However, 29 out of 113 crabs measured over three days were found on sediment that was 40°C or hotter, and seven of these individuals had *T*_*b*_s that were 40°C and higher. All of these individuals were active and appeared healthy although we have no information on the frequency or duration of these periods of very high *T*_*b*_ and their impact on crab health and survival. If we had collected data during the hottest month of the year (July), it is probable that the median *T*_*b*_s and the frequency of very hot crabs would have been higher.

The linear relationships between thermoregulation capacity (*T*_*b*_ − *S*) and surface temperature that we found for females of all sizes and for small males allowed us to explore the impact of warming surfaces on crab bodies. For females, we found that for every 1°C of surface warming, their bodies warmed about 0.7°C, a finding that quantifies the degree to which they resisted heat transfer from the surface. Quantifying changes in behavior, especially with respect to use of burrows and foraging locations, along with an energy budget analysis, could help to elucidate the costs of this resistance. Females in both size categories were large enough to mate and carry broods, so we did not expect behavioral differences related to reproductive behaviors or embryo incubation to impact their responses to surface warming. Larger females tended to be around a third of a degree cooler than smaller ones on surfaces of the same temperature, a difference that might have been due to a higher thermal inertia or longer legs lifting them higher above the substratum. The bodies of small males warmed about 0.77 °C for every 1°C of surface warming, but we do not know if this represents a significant difference from the other groups because we did not compare small and large males or males to females.

Large males were in a class of their own, with far less of the variation in thermoregulation capacity explained by surface temperature compared to females and small males. There was a non-linear relationship between thermoregulation capacity and surface temperature, with some large males much cooler relative to the surface than expected for a linear relationship. It is not surprising that the relationship between thermoregulation capacity and surface temperature would be different for this group. Male fiddler crabs experience allometric growth during development, with the large claw becoming disproportionately larger as the crab grows ([44]; [45]; [46]), and this sexually selected appendage is known to also function as a heat sink that lowers core body temperatures ([20]). Furthermore, claw and body sizes influence abilities to attract mates and defend burrows ([47]; [31]; [48]), and courtship behaviors impact thermoregulation. Males that are actively courting females may experience high body temperatures if they are displaying in open habitats where they can be seen or higher in the intertidal where females prefer the more stable burrows ([49]; [50]). However, large courting males also crowd into shaded areas that give them a thermal advantage ([23]; [51]; [52]), and those actively defending burrows have ready access to a cool microclimate. Hence, individual courting males may experience higher or lower than expected *T*_*b*_s depending on local conditions, including the availability of shade, female habitat preferences, and variation in the intensity of sexual competition. Because we collected crabs opportunistically, we probably collected some large males that were actively courting and some that were not. Although it is not clear to us whether courting or non-courting males would have been expected to have cooler *T*_*b*_s at the same surface temperatures, sorting large males into different behavior categories might have elucidated the relationship between their thermoregulation capacity and surface temperature. Alternatively, this relationship might simply be inherently more variable and nonlinear for this group.

Fiddler crab habitats are riddled with burrows that extend tens of centimeters into the substratum, providing refuge from predators and inhospitable surface temperatures. Males entice females into burrows for matings and some species brood in them. The amount of time spent in burrows is influenced by season and local environmental conditions. During the daytime low tide exposure period, *M. panacea* spends around a quarter of its time in a burrow during the non-breeding season but nearly half of its time there while breeding ([24]). The fiddler crab *Austruca mjoebergi* is almost as likely to be in its burrow as it is feeding on the surface when its burrow is in the sun, but time in the burrow drops dramatically in shade, where it is more than three times as likely to be feeding on the surface ([52]). The fact that crabs spend less time in burrows when those burrows are shaded suggests a costly tradeoff between feeding and thermoregulation, a cost that may increase for *M. pugnax* during the reproductive season when it experiences declines in fat stores ([53]). During 2019 on Sapelo Island, Georgia, USA burrows offered an important, if potentially costly, thermal refuge for *M. pugnax*. We found that most of the thermoregulation capacity for females was explained by the cooling capacity of the burrow–as the burrow became cooler relative to the surface, so did female *T*_*b*_. This was especially true for large females where over 70% of their thermoregulation capacity was explained by the cooling capacity of the burrow. We found that for a given cooling capacity, larger females were cooler than smaller ones when burrow and surface temperatures were held constant, and large females tended to be cooler than small ones when the burrow could not be used for thermoregulation (i.e., when burrow and surface temperatures were the same). These consistent findings could be explained by thermal inertia and the longer legs of large females. We estimated that for every 1.0 °C degree cooler the burrow became relative to the surface, large females could cool their bodies down an additional 0.5°C.

Males clearly used the burrow less efficiently for thermoregulation, cooling their bodies by around a third of a degree for every 1°C increase in the cooling capacity of the burrow. They were also only around a degree hotter than the surface when the burrow could not be used for thermoregulation, compared to small females which were 2°C hotter. Furthermore, less of the variation in thermoregulation capacity was explained by the cooling capacity of the burrow, with only about 50% of that variation explained for large males. Males appear to rely less on the burrow for thermoregulation than females, quite possibly because their enlarged claw provides an additional means for cooling down. They are also larger, overall, which gives them a thermal advantage when the surface heats up. Our estimates of the burrow use efficiency (*E*_*B*_) are conservative because we used burrow temperatures at 30 cm in our calculations, a reference depth that captured the coolest burrow microclimate available. If crabs were actually retreating to shallower depths where temperatures were not as cool, then the cooling capacity would have been reduced and the burrow use efficiency (*E*_*B*_) estimate would have been higher. In future work, measurements of how deep crabs go and their duration at these depths could be used to refine estimates of *E*_*B*_. We also think that it would be important to compare the cooling capacity of burrows in different habitats and climates, as places with similar surface and air temperatures but different burrow cooling capacities would present dissimilar thermal landscapes. For example, it appears that *M. pugnax* in the southern United States experiences surface temperatures in the summer that are similar to those experienced by *Tubuca urvillei* in Kenya. However, the surface and 20 cm depth temperatures measured by Fusi et al. (2015) in Kenya were not very different, suggesting that the cooling capacities of burrows there may be low. If this is true, it would not be surprising to find that *T. urvillei’s* thermoregulation capacity is lower than that of *M. pugnax*, which would result in different evolutionary pressures and responses to climate warming for the two species.

Finally, we found that male and female crabs of different sizes showed different thermoregulation capacities, burrow use efficiencies, and *T*_*b*_s when the burrow could not be used for thermoregulation. However, the *T*_*b*_ at which they all started using burrows to thermoregulate (*T*_*reg*_) was around of 24°C, suggesting that *T*_*reg*_ is a parameter that is set by *M. pugnax’s* physiology, regardless of sex and size. Because we used data collected from March through October, this *T*_*reg*_ estimate should be viewed as an average for the 2019 breeding season on Sapelo Island, Georgia. In future work, seasonal and latitudinal differences in *T*_*reg*_ could be investigated to determine the degree to which acclimation and local adaptation influence this parameter. We also think that questions about the relationships between *T*_*reg*_, *T*_*o*_, and *T*_*set*_ should be investigated. Is the *T*_*b*_ at which field-based crabs start to use burrows (*T*_*reg*_) closer to the preferred *T*_*b*_ of *M. pugnax* in the lab (*T*_*set*_) or to the optimum temperature for some aspect of its performance (*T*_*o*_)? If *T*_*reg*_ and *T*_*set*_ are different, do crabs start to use burrows before they reach *T*_*set*_ or after they’ve passed it? Answering these questions will elucidate tradeoffs and strategies used by crabs as they respond to their warming environments.

## Conclusion

While very hot days at the peak of summer can push intertidal invertebrates close to or above their *CT*_*max*_, the deleterious impacts of warmer days earlier in the reproductive season are less obvious. For *M. pugnax*, changes in burrow use related to thermoregulation involve fitness trade offs because time spent in burrows results in lost feeding and courtship opportunities ([54]; [55]; [30]). However, while using a burrow to thermoregulate is potentially costly, it is also highly effective. As the surface warms rapidly through the day during the reproductive season in temperate climates, burrow temperatures remain cool and stable. For example, during a 48 hour period in May, we found temperatures in artificial burrows that were nearly 20°C cooler than the surface in the mid-afternoon when surface temperatures were hottest (burrows were warmer than the surface from evening to late morning; unpublished data). We found that crabs used burrows to counteract *T*_*b*_ increases, and we estimated *T*_*reg*_, the *T*_*b*_s at which crabs began to use burrows for thermoregulation, to start at around 24°C for *M. pugnax*. In late March, we found that many crabs showed *T*_*b*_s at or above this temperature, and when we returned in May the median *T*_*b*_s for crabs of all sizes and both sexes were above *T*_*reg*_. This was still the case for our last collection in October. With climate warming, *M. pugnax* will experience higher metabolic rates like all ectotherms, but there will be additional energy budget impacts as crabs spend more time cooling down in burrows beginning earlier in the year.

